# Sex-specific differences in ADAR editing patterns in response to SARS-CoV-2 infections

**DOI:** 10.64898/2026.09.29.755432

**Authors:** Aiswarya Mukundan Nair, Helen Piontkivska

## Abstract

Sex-specific differences in COVID-19 outcomes are well documented, with male sex being a key risk factor for severe disease. Multiple factors, including differences in the expression of Interferons (IFNs) and Interferon stimulated genes (ISGs) between the sexes have been identified as contributing to these differences. Adenosine-to-inosine (A-to-I) RNA editing, mediated by ADAR enzymes, is an important component of antiviral innate immune signaling. While previous studies, including us, have identified altered expression and activity of ADARs in response to SARS-CoV-2 infection, it remains unknown whether sex-specific differences in expression and activity of ADARs exist during infection with SARS-CoV-2. We address this question by analyzing a publicly available whole blood RNA sequencing dataset from males and females, matched for age and comorbidity status across pre-, mid-, and post-infection stages of SARS-CoV-2 infection. Our results show distinct patterns of ADAR expression between the sexes across different stages of infection, with significant differences in the expression of ADAR and ADAR2 in pre- and mid-infection that were not observed post-viral clearance. Interestingly, expression of predominantly neural isoform ADAR3 was significantly different and higher in females, mid-infection. Furthermore, these differences in ADAR expression were likely amplified downstream with significant differences in the global editing levels. Correlation analysis revealed that the relationship between ADAR expression and global editing activity differed by sex and stage of infection, indicating that ADAR editing is dynamically regulated and is influenced by factors other than ADAR transcript abundance. Noteworthy, sex-specific differences in ADAR editing were observed within genes that escape X-chromosome inactivation (XCI), a gene set enriched in immune related functions and known to be involved in sex-specific outcomes to viral infections. ADAR editing within these genes differed at baseline and changed across stages of infection in a manner independent of gene expression. Altogether, our findings establish ADAR editing as an additional layer of immune regulation that may be contributing, in part, to the differential clinical manifestations of COVID-19 observed between males and females.

## Introduction

Differences in susceptibility, immune responses, and outcomes to both DNA or RNA virus infections is widely observed between males and females. Although the contribution of age and other host factors cannot be excluded, a general trend of lower immune responses and higher adverse effects is consistently observed in males (Dudley & Lee, 2020; Gay et al., 2021; Scully et al., 2020). For instance, males are more prone to higher incidence and/or severe disease outcomes in response to infection with Epstein-Barr virus, hepatitis B (HBV), hepatitis C (HCV), seasonal and pandemic influenzae A virus (H1N1) (Choi et al., 2011), Herpes Simplex Virus (HSV-1) (Torcia et al., 2012), rhinoviruses (Carroll et al., 2010), and West Nile Virus (WNV) (Hoffman et al., 2019). Similar sex-specific differences are also observed in infection caused by betacoronaviruses, including MERS (Alghamdi et al., 2014), SARS (Karlberg et al., 2004; Leong et al., 2006), and SARS-CoV-2. During the COVID-19 pandemic it was observed that despite similar rates of infection between males and females (Bhattacharya et al., 2024; Viveiros et al., 2021), males developed more severe manifestations and were at a higher risk of death from SARS-CoV-2 infections, compared to matched females (Marina & Piemonti, 2020; Scully et al., 2020). Moreover, the symptoms observed between the two sexes were mostly similar, although differences were observed in the severity of specific symptoms (Massion et al., 2023; Sauerwald et al., 2022). These differential responses arise from a combination of factors, including immunomodulatory functions of sex hormones, variability in the expression of X chromosome-encoded genes and host entry receptors, immune related differences, and sex-dependent host-virus interactions (Bongen et al., 2019; Márquez et al., 2020; Spiering & de Vries, 2021).

High levels of pro-inflammatory cytokines are associated with severe COVID-19 pathology in males (Del Valle et al., 2020; Gustine & Jones, 2021; Merad & Martin, 2020; Queiroz et al., 2022). Studies in animal models have shown that the predominantly female sex hormone, estrogen (especially in pre-menopausal women), inhibits the expression of inflammatory cytokines such as IL-1, IL-6, and TNF-α; this anti-inflammatory/immunomodulatory role of estrogen is suggested to have protective effects in females during SARS-CoV-2 infections (Al-Lami et al., 2020a; Gaskins et al., 2012; Straub, 2007; Takehiro & Akiko, 2021). Although encoded on the X chromosome, the expression of SARS-CoV-2 entry receptor angiotensin converting enzyme 2 (ACE2) is downregulated by estrogen. This hormone-mediated reduction in ACE2 expression is suggested to be protective against severe COVID-19, similar to the milder outcomes observed in children, who also show lower ACE2 expression. In contrast, expression of transmembrane serine protease 2 (TMPRSS2) - that primes viral entry into the cell - is regulated by male androgens and has been implicated in more severe outcomes in males (Al-Lami et al., 2020b; Bach & Jensen, 2025; Bunders & Altfeld, 2020; Koyou et al., 2025). Furthermore, multiple immunoregulatory genes (e.g., TLR7, TLR8) and those essential for virus-host interactions (e.g., ACE2) are encoded on the X chromosome. Many of these genes escape X chromosome inactivation (XCI) in females, which is meant to maintain gene dosage between the sexes (Tukiainen et al., 2017; Youness et al., 2023; Souyris et al., 2018; Spiering & de Vries, 2021). Escape from XCI results in higher expression of these immune-related genes and a more robust immune response in females. For instance, TLR7, one of the key pattern recognition receptors for SARS-CoV-2 ssRNA, escapes XCI, resulting in its higher expression in females. This biallelic expression of TLR7 enables females to mount a stronger and more protective immune response during SARS-CoV-2 infections (Gómez-Carballa et al., 2022; Keur et al., 2024). This difference in the dosage of TLRs and other immune pathway components is reflected downstream in the expression of its effectors, including interferon (IFNs) and interferon stimulated genes (ISGs). For instance, TLR8 can evade XCI contributing to higher TLR8 protein levels in immune cells, affecting response to infection in females (Youness et al., 2023). Notably, elevated TLR7 expression at mRNA and protein levels leads to stronger TLR-7 mediated IFN and ISGs expression in females during SARS-CoV-2 infection (Hagen et al., 2020; Souyris et al., 2018; Spiering & de Vries, 2021).

Heterogeneity in the expression of IFNs and ISGs is thought to be a key factor contributing to severe outcomes in COVID-19 (Gozman et al., 2021; Hadjadj et al., 2020; Li et al., 2021; Masood et al., 2021; Scagnolari et al., 2021). Multiple studies have established sex-specific differences in the expression of IFNs and ISGs in response to SARS-CoV-2 infections, with females in general showing higher levels than males (Agrawal et al., 2021; Takahashi et al., 2020). Many such ISGs, in addition to their potential antiviral effects, have other roles essential for normal physiological functions. One such ISG is an isoform of Adenosine Deaminases that Act on RNA (ADAR), specifically the ADAR1 isoform ADARp150, which is a member of ADAR protein family (Eisenberg & Levanon, 2018). These proteins are encoded by three genes within humans, of which ADAR1 and 2 encode catalytically active enzymes, while the third ADAR3 has no documented editing activity (Raghava Kurup et al., 2022; Y. Wang et al., 2019). ADARs are post-transcriptional RNA editing enzymes that deaminate adenosines (A) to inosines (I) within double stranded RNAs (dsRNAs) substrates. ADAR1 also exists as another isoform, ADARp110, whose expression also appears to be elevated during viral infections (e.g., Nachmani et al., 2014; Mukundan Nair & Piontkivska, 2026a). Because inosine is interpreted as guanosine (G) by the translational machinery, ADAR edits can have far-reaching functional consequences by altering coding sequences, changing protein function, affecting splicing, miRNA binding and targeting and/or serving as a mechanism to regulate gene expression and transcript diversity (Liu et al., 2024; Rehwinkel & Mehdipour, 2025; Samuel, 2011). In addition to its transcriptome regulation role, ADARs are key components of antiviral innate responses through their pro- or anti-viral effects (Piontkivska et al., 2021). While viral infections mostly alter expression of ADAR1, in turn, ADAR1 can interact with other ADARs and change their activity and editing within their substrates (Lamers et al., 2019; Quin et al., 2021). Therefore, an increase in ADAR1 levels could affect transcriptome-wide editing. ADAR editing within transcripts is highly nuanced (Huntley et al., 2016; Tariq & Piontkivska, 2024; Mercer et al., 2025; Mukundan Nair & Piontkivska, 2026a) and is precisely maintained during normal physiology. However, irregularities in editing are observed during multiple disease states, including neurological disorders (Sodhi et al., 2024; Yang et al., 2021), cardiovascular disorders (Altaf et al., 2019), metabolic disorders (Kung et al., 2024), cancers (Baker & Slack, 2022), and autoimmune conditions (Magni & Ceruti, 2020). Moreover, ADARs role as a mechanistic link between viral infections and both immediate and long-term clinical manifestations is increasingly being appreciated (Piontkivska et al., 2019; Piontkivska et al., 2021).

Previous studies have identified widespread ADAR editing signatures within SARS-CoV-2 genome. Host ADAR mediated editing in this scenario is known to impact viral fitness, transmissibility, and evolution (Di Giorgio et al., 2020; Picardi et al., 2021). Of particular note are the non-synonymous ADAR edits within the receptor binding domain of viral spike protein that can potentially alter viral infectivity (Ringlander et al., 2022). Furthermore, redirection of some of the editing activity from the virus to host transcriptome is also reported, where editing is proposed to contribute to acute symptoms (Jin et al., 2024), varying disease severity (Mukundan Nair & Piontkivska, 2026a), and to post-acute sequelae of COVID-19 (PASC) (Mukundan Nair & Piontkivska, 2026b). These studies establish the importance of dynamic ADAR editing in COVID-19 disease manifestations. To the best of our knowledge no studies have identified whether ADAR editing signatures could differ between sexes during viral infections, potentially contributing to the sex-specific outcomes prevalent during viral infections. However, it should be noted that sex has been identified as a variable affecting expression of ADAR1 and ADAR2 (Giacopuzzi et al., 2018), and sex-specific ADAR editing signatures have been identified in multiple diseases. For example, altered ADAR editing that differs in a sex-specific manner holds prognostic potential, and is associated with survival, in glioblastomas and isocitrate dehydrogenase mutated (IDH-MUT) gliomas (Lin & Chen, 2022). Similarly, sex-specific ADAR editing signatures have been observed in major depressive disorders (MDD), Parkinson’s disease, Alzheimer’s disease (Gupta et al., 2025), and during inflammatory states (Mercer et al., 2025; Nichols et al., 2022), indicating that ADARs can act in a sex-specific manner during disease states. Therefore, taking SARS-CoV-2 as a model, here we examine whether expression and activity of ADARs could differ between males and females. To address this, we first analyzed ADAR expression and activity pre-infection to establish potential baseline differences, followed by analysis during active infection, and post viral clearance, using whole blood sequencing data from males and females with mild COVID-19 that were sampled pre-, mid-, and post-stages of infection (GSE198449) (Sauerwald et al., 2022).

## Methods

### RNA sequencing dataset

ADAR expression and editing profiles across three distinct stages of SARS-CoV-2 infection, of age-matched males and females who developed mild COVID-19, were investigated from the publicly available whole blood RNA sequencing data from Sauerwald et al. study (BioProject PRJNA815324; Sauerwald et al., 2022). The dataset included samples collected from young marine recruits enrolled in the COVID-19 Health Action Response for Marines (CHARM), and were sampled pre-, mid- and post-SARS-CoV-2 infection as a part of tracking disease spread. We selected only those samples that were reported to have developed mild COVID-19, with symptoms including abdominal pain, chills, cough, decreased sense of taste or smell, diarrhea, fatigue, fever, headache, muscle ache, nausea/vomit, runny nose, shortness of breath, and sore throat, and where infections were confirmed using quantitative PCR of nasal swabs. To reduce disparities in sample size, while including a maximum number of samples per group, we randomly selected a comparable number of samples per group. This resulted in 89 female and 100 male samples pre-infection, 89 female and 100 males mid-infection, and 56 females and 100 males post-infection for the present study (Table 1).

**Table 1:** Characteristics of subset of CHARM dataset (BioProject PRJNA815324) used in this study.

|  | **Pre-infection** |  | **Mid-infection** |  | **Post-infection** |  |
| --- | --- | --- | --- | --- | --- | --- |
| Sex | Males | Females | Males | Females | Males | Females |
| Total number of samples | 100 | 89 | 100 | 89 | 100 | 56 |
| Average number of mapped reads<br>(approximate value in million mapped reads/sample) | 22.0 | 27.4 | 25.6 | 21.2 | 24.0 | 24.0 |
| Mean number of ADAR editing events per sample for each group | 485661 | 565512 | 586560 | 503416 | 544616 | 520951 |

### Computational analysis of RNA sequencing data and variant calling to identify ADAR edits

Analysis of RNA sequencing data was performed, as described before, using Automated Isoform Diversity Detector (AIDD) (Plonski et al., 2020; Mukundan Nair & Piontkivska, 2026b), a computational pipeline that uses various open-source tools for raw RNA sequencing analysis and variant calling. To elaborate, raw sequencing reads publicly available at NCBI-SRA under GEO accession GSE198449 (PRJNA815324) (Sauerwald et al., 2022) were downloaded and checked for sequence quality using FASTQC (https://www.bioinformatics.babraham.ac.uk/projects/fastqc/). The alignment program HISAT2 (Kim et al., 2015), which provides fast as well as sensitive mapping, was then used to align reads to human reference genome GRCh37. Stringtie (Pertea et al., 2015) was then used for transcriptome assembly and to quantify mapped reads, including the expression levels of ADARs in Transcripts Per Million (TPM). Analysis with DESeq2 (Love et al., 2014) followed pairwise design comparing gene expression between males and females at each stage of infection. Variant calling was done using Java framework for accurate SNV assessment (JACUSA2) (Piechotta et al., 2022), an analysis framework used to identify post-transcriptional RNA modifications from the aligned BAM files. A filter to exclude sites with less than 20 reads per site was used to identify putative ADAR editing sites and count the number of bases observed at each such site. Both A-to-G and U(T)-to-C substitutions were defined as ADAR editing sites, the latter representing editing sites on the complementary strand. The output matrixes were annotated, including for genomic location, genes incorporating the edits, and type of substitution using ANNOVAR gene symbols using RefGene (K. Wang et al., 2010) and SnpEff (Cingolani et al., 2012) to get the predicted impact of these edits.

### Calculation of per site editing rate and global ADAR editing rates

JACUSA2-generated VCF files were filtered to include only those A-to-G and T-to-C substitutions with total number of aligned reads ≥ 100. From the remaining sites, only those sites that were previously defined in REDIportal database (Mansi et al., 2020) were included. REDIportal is a database containing millions of pre-identified, annotated ADAR editing sites. Filtering our list to include only sites present in REDIportal was done to increase the confidence of selected edited site and to reduce potential false positives. Per site editing rates of the filtered sites were calculated by dividing the total number of edited reads at a site (total number of G’s if the reference allele is A, and total number of C’s if the reference is T) to the total number of reads aligned to that site. Overall ADAR editing levels, calculated as unique value per sample that represent the global levels of editing within a sample, were computed using methods defined in Breen et al 2019 (Breen et al., 2019). Briefly, for each sample, the overall ADAR editing levels were calculated as the ratio of total number of edited reads to the total number of all reads aligned to ADAR editing sites identified in that sample. Difference between the overall ADAR editing rates between sexes were assessed using Wilcoxon signed-rank test. Differences within sexes, across the stages of infection, were assessed using ANOVA.

### Statistical analysis of ADAR expression and correlations with global editing

To understand the relationship between ADAR expression and the overall ADAR editing levels, Spearman correlation coefficients (*ρ*) that represents the strength and direction of monotonic relationships were calculated between the expression levels of all three ADARs measured in TPM and the overall ADAR editing levels. Analyses were conducted separately for each sex and stage of infection to understand the relationship for within a sex for each stage of infection for all ADAR isoforms. Coefficient of determination (R^2^) that represents the proportion of variance in the overall ADAR editing that could be explained by the expression of ADARs, were derived from linear regression model.

### List of genes escaping X chromosome inactivation

Curated list of X chromosome inactivation (XCI) escape genes were obtained from Tukiainen et al. 2017 (Tukiainen et al., 2017) (Supplementary Table 3). Gene symbols from this list was matched to SnpEff gene annotations. ADAR editing sites mapping to genes that escape XCI were retained for X chromosome specific analyses. For each sample and each XCI escape gene weighted editing levels were computed. Weighted editing levels are defined as the proportion of edited reads across all edited sites within a gene which is calculated as the sum of edited reads divided by sum of total reads across all edited sites. Differences in weighted editing levels between sexes were assessed using Wilcoxon rank-sum test. Effect sizes were calculated using Cliff’s delta.

## Results

### Differences in expression of ADAR genes

To examine whether levels of ADARs differs by sex and to track temporal changes in their expression within each sex, we compared ADAR expression, measured as Transcripts Per Million (TPM), between the two sexes at each stage of infection as well as within each sex across all three (pre-, mid-, and post-) stages of SARS-CoV-2 infection. ADAR1 expression levels differed significantly between the sexes, with higher expression in females, both pre- (t-test: p-value = 2.7e-05) and mid-infection (t-test: p-value = 0.0009), although this difference was not observed post-infection (t-test: p-value = 0.3580). Similar pattern of significant differences in ADAR1 expression were observed within both males (ANOVA: p-value = 4.79e-08) and females (ANOVA: p-value = 5.44e-15) across all three infection stages (Fig 1A). Likewise, ADAR2 expression levels displayed significant differences between males and females pre- (t-test: p-value = 0.0013) and mid- (t-test: p-value =0.0046) infection stages, with lower expression in pre- and higher expression in mid-stages of infection in females, respectively. As was observed for ADAR1, ADAR2 levels also displayed no significant differences between sexes post-infection (t-test: p-value = 0.3999). Notably, significant differences in ADAR2 expression across all three stages were observed only in females (ANOVA: p-value = 5.3e-12), while the difference in ADAR2 expression across all three stages within males were not significant (ANOVA: p-value = 0.129) (Fig 1B). Interestingly, expression of a predominantly neural gene, ADAR3, while not significantly different pre- (t-test: p-value = 0.7237) and post-( t-test: p-value = 0.3737) infection between sexes, displayed significantly higher levels in females mid-infection (t-test: p-value = 0.0073), and was also found to differ significantly across all three stages within females (ANOVA: p-value = 0.0294), but not males (ANOVA: p-value = 0.6) (Fig 1C).

**Figure 1:**
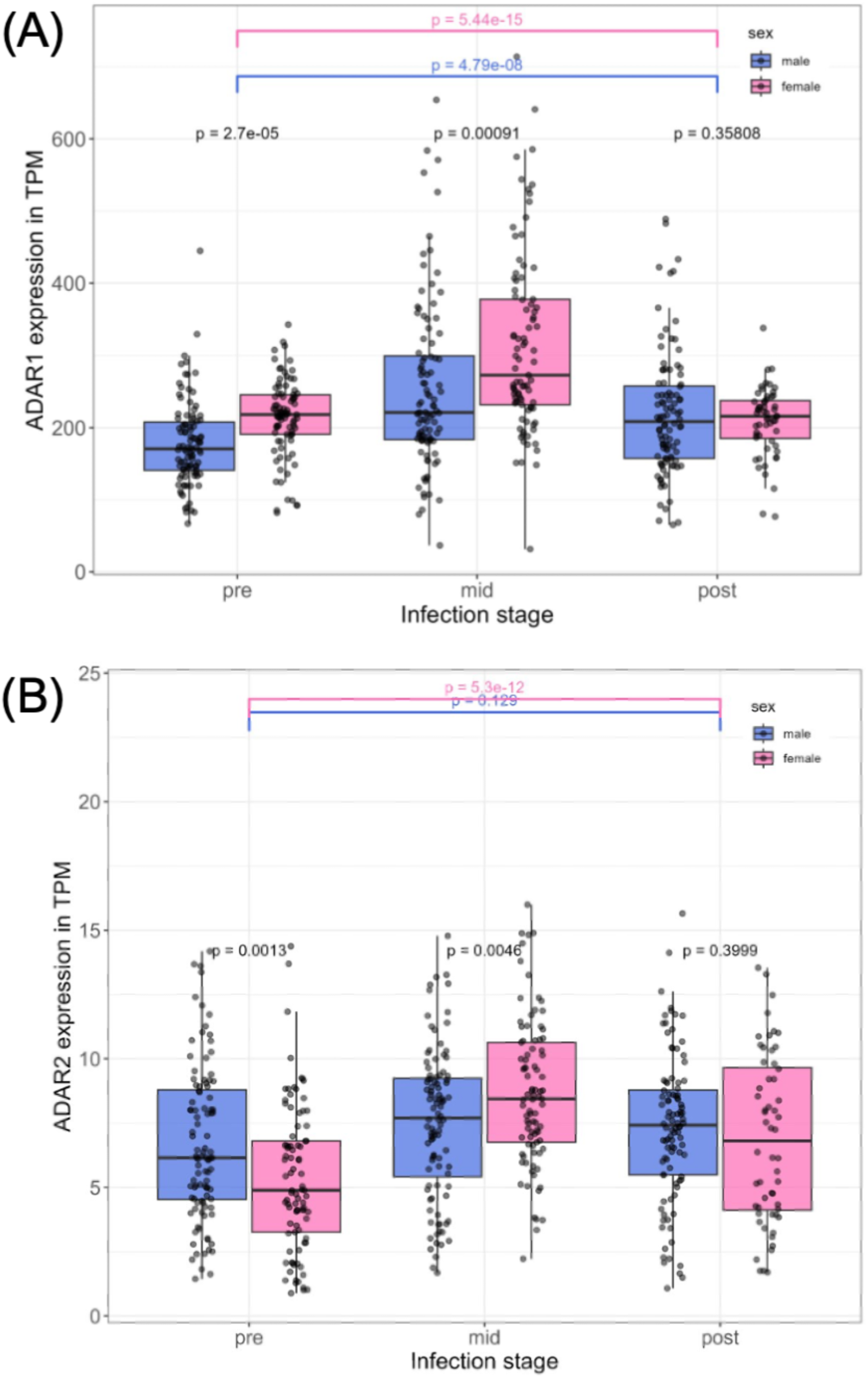
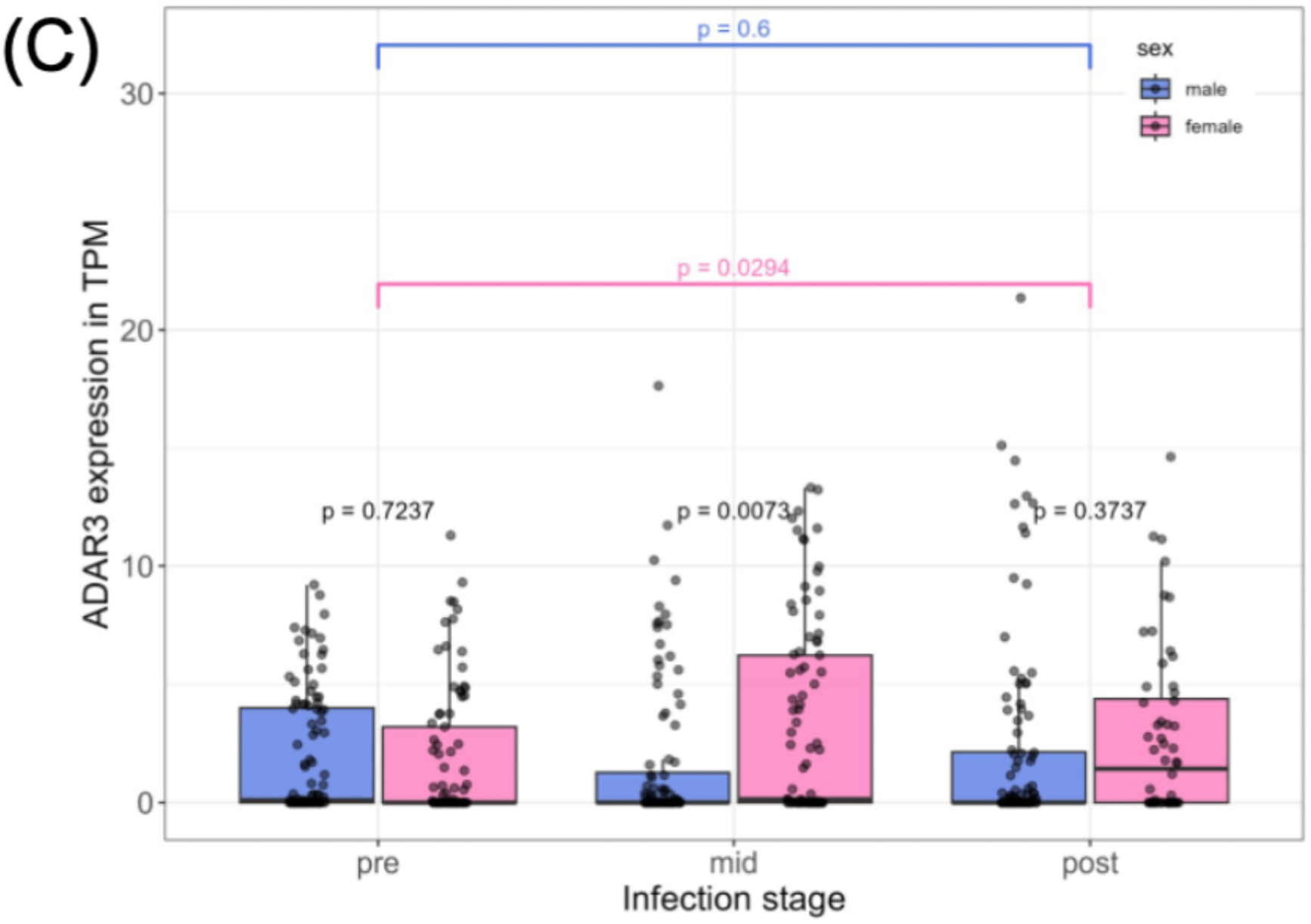
Difference in expression of ADAR genes between sex at three distinct stages of infection and within each sex across all three stages of infection. (A) ADAR1 expression is significantly higher in females both pre- (t-test: p-value = 2.7e-05) and mid-infection (t-test: p-value = 0.0009) and are significantly different within males (ANOVA: p-value = 4.79e-08) as well as females (ANOVA: p-value = 5.44e-15) across all three stages of infection. (B) Pre-infection levels of ADAR2 are significantly lower in females (t-test: p-value = 0.0013). While mid-infection females show higher levels of ADAR2 (t-test: p-value =0.0046) and significant differences in ADAR2, across the three stages is observed only in females (ANOVA: p-value = 5.3e-12) but not males (ANOVA: p-value = 0.129). (C) ADAR3 expression showed significant differences only mid-infection (t-test: p-value = 0.0073) and in females (ANOVA: p-value = 0.0294) across all three stages of infection, while males had negligible expression across all three stages of infection (ANOVA: p-value = 0.6).

### Global changes in ADAR editing between the sexes across the three stages of infection

Considering differences in ADAR expression between sexes that changed across all three stages of infection, we next sought to investigate quantitative differences in ADAR editing. To address this, we measured a) difference in the total number of ADAR edits within each group, and b) differences in the overall ADAR editing levels, defined as the ratio of all the edited reads from REDIportal-identified ADAR edited sites in a sample to the total number of reads aligning to these sites. This analysis was done between sexes across each stage of infection, and within each sex across all stages of infection. Among all the identified substitutions, A-to-G and T-to-C representing putative ADAR edited sites were most abundant across all three stages in both sexes (Fig 2A), although these substitutions displayed differences between sexes across all three stages (Fig 2B). Females had an average of 565,512 ADAR edits per sample (Supplementary Table 1A), which was higher than males, who had an average of 485,661 ADAR edits per sample pre-infection (Supplementary Table 1B). In mid-infection the average number of ADAR edits was higher in males (586,560 edits/sample) (Supplementary Table 1D) compared to females (503,416 edits/sample) (Supplementary Table 1C). Post-infection, the average number of ADAR edits remained higher in males (544,616 edits/sample) (Supplementary Table 1F) compared to females (520,951 edits/sample) (Supplementary Table 1E). Furthermore, the overall ADAR editing levels, representing editing rates per sample, did not differ significantly at baseline (pre-infection) between sexes (Wilcoxon rank-sum test: p-value = 0.6516) (Supplementary Table 2 A, B). Notably, editing rates differed significantly between males and females both mid- (Wilcoxon rank-sum test: p-value = 0.0055) (Supplementary Table 2 C, D) and post-infection (Wilcoxon rank-sum test: p-value = 0.0297) (Supplementary Table 2 E, F). However, within sex, no significant difference was observed in the overall ADAR editing rates across all stages (Fig 2C).

**Figure 2:**
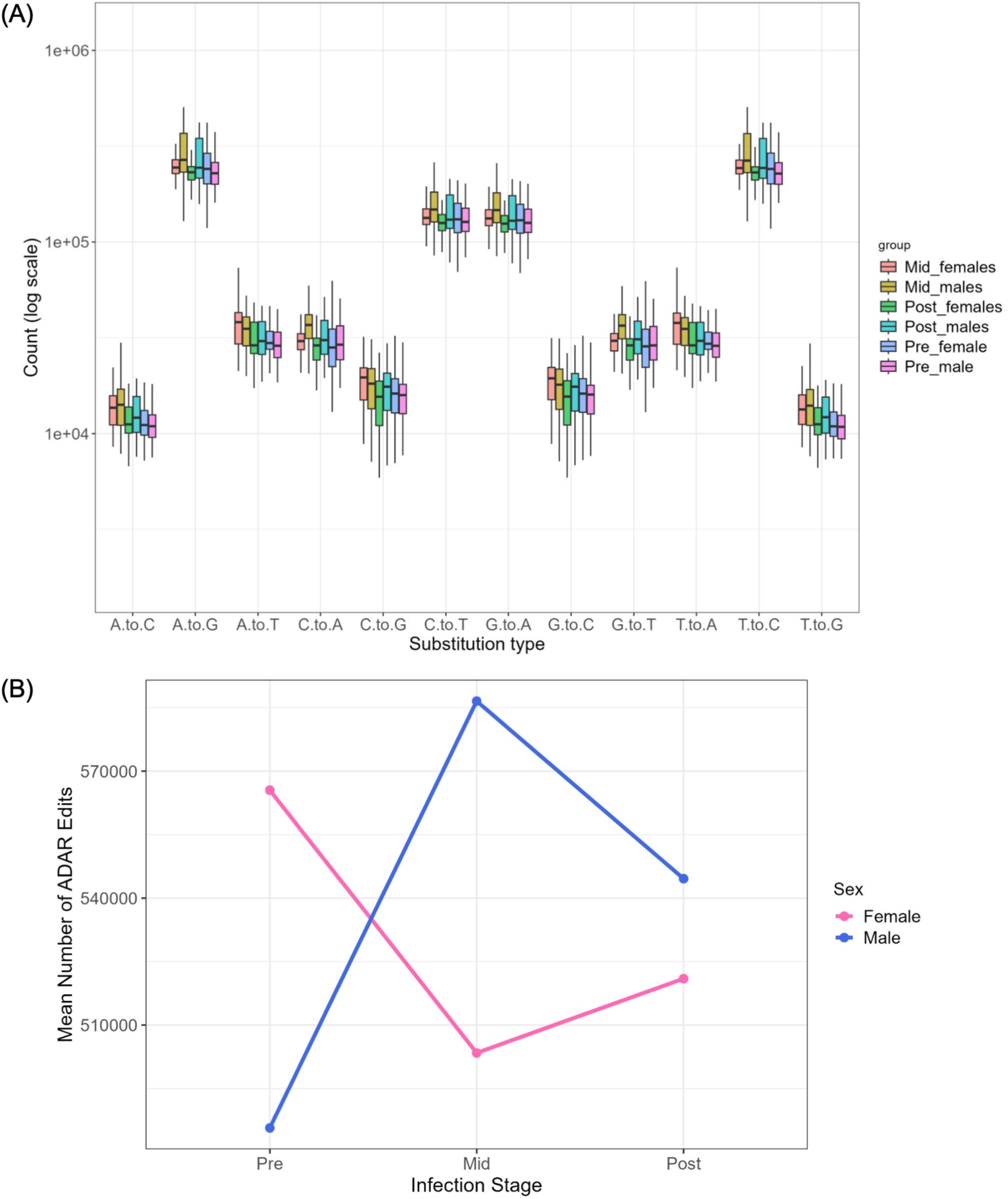
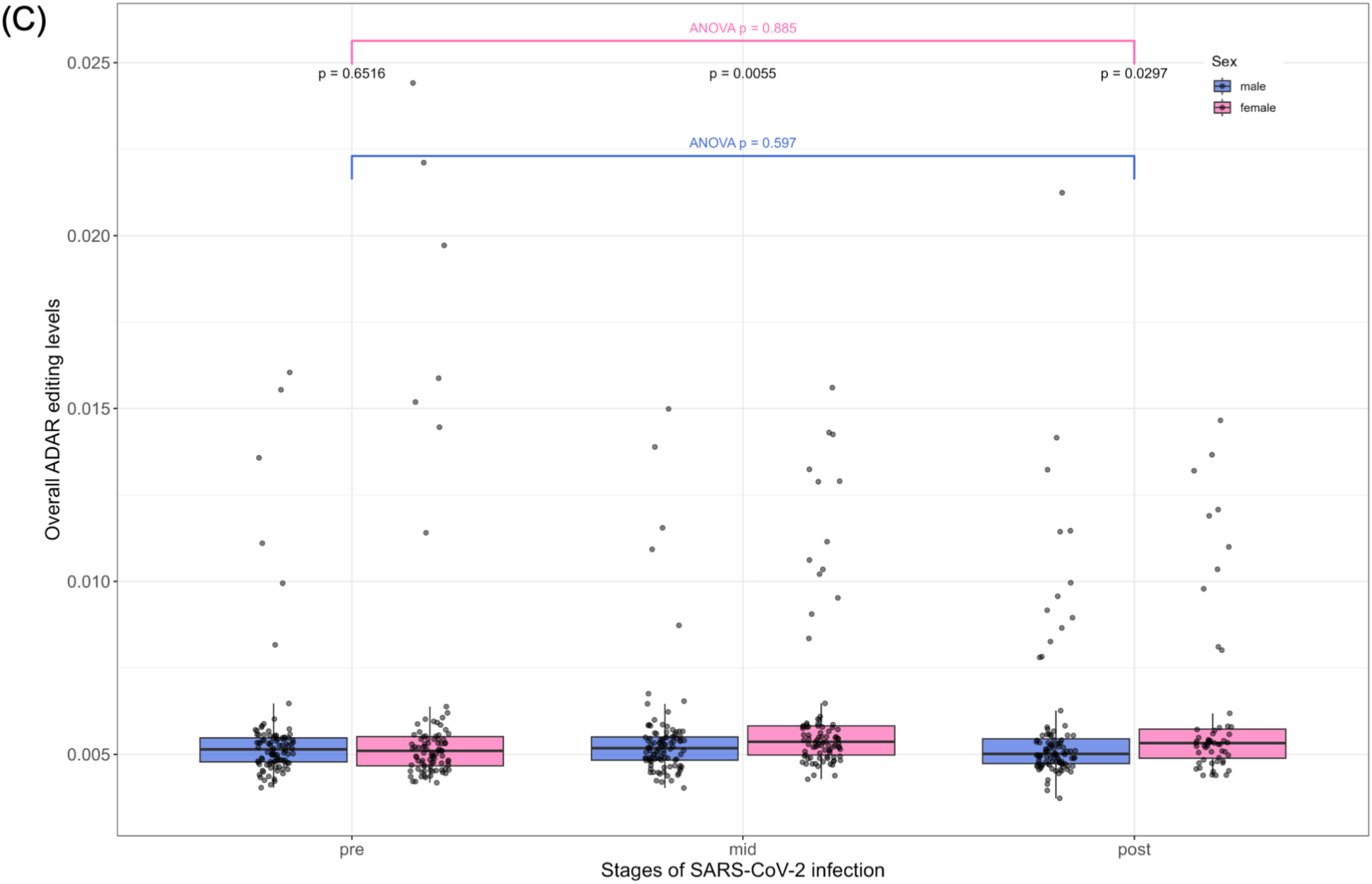
Transcriptome wide changes in ADAR editing between sexes across pre-, mid-, and post-stages of infection. (A) Distribution of various genomic substitutions including A-to-G and T-to-C representing potential ADAR edited sites, across all three stages of infection in males and females. (B) Average number of ADAR edited sites per sample between males and females across the three stages. (C) Differing overall ADAR editing levels between sexes across all three stages and within sex across the stages.

### Correlations between expression levels of ADARs and editing activity

Next, to examine whether the observed differences in the overall ADAR editing levels were driven by expression of ADARs, we calculated correlations between the expression values of all three ADARs measured in TPMs with the overall ADAR editing levels within each sex across all three stages of infection. Significant positive correlation between ADAR1 expression and the overall ADAR editing was found only in males during mid-infection (*ρ =* 0.28, R^2^ = 0.076, p = 0.0056). It was slightly positive pre-infection (*ρ* = 0.012, p > 0.05) and slightly negative post-infection (*ρ =* -0.07, R^2^ = 0.005, p >0.05). While in females, this correlation - though not statistically significant - was negative across all stages (pre -infection: *ρ =* -0.13, R^2^ = 0.016, p >0.05; post-infection: *ρ =* -0.22, R^2^ = 0.05, p >0.05), including mid-infection (*ρ =* -0.16, R^2^ = 0.026, p >0.05) (Fig 3A), suggesting sex-dependent ADAR1 activity during active infection. In contrast to ADAR1, ADAR2 expression displayed a significant positive correlation with the overall ADAR editing levels mid-infection in both males (*ρ =* 0.29, R^2^ = 0.084, p = 0.0035) and females (*ρ =* 0.028, R^2^ = 0.08, p = 0.0073), while no significant correlations were found between ADAR2 expression and editing levels in other comparison groups (Fig 3B). This suggests that the overall ADAR editing levels within males might be driven by the expression of both ADAR1 and ADAR2, while ADAR2 might be the primary driver of changes in the overall ADAR editing levels mid-infection in females. Furthermore, no significant correlation was observed between ADAR3 expression and the overall ADAR editing in any of the comparison groups (Fig 3C), which can potentially be attributed to the lack of enzymatic activity of ADAR3. Notably, although the magnitudes of significant correlation coefficients for ADAR1 and ADAR2 show fair associations (*ρ* < |0.3|) (Akoglu, 2018), their statistical significance suggests biological relevance and the sex- and enzyme-specific regulation of the overall ADAR editing across distinct stages of infection. These results indicate that the ADAR activity might differ between sexes across the three stages of SARS-CoV-2 infection, and that factors other than ADARs might also be contributing to differences/changes in the overall ADAR editing levels. Furthermore, while the remaining comparison did not reach statistical significance, the correlation coefficients provide directional relationships between ADAR expression and overall editing (Akoglu, 2018) for each comparison group.

**Figure 3:**
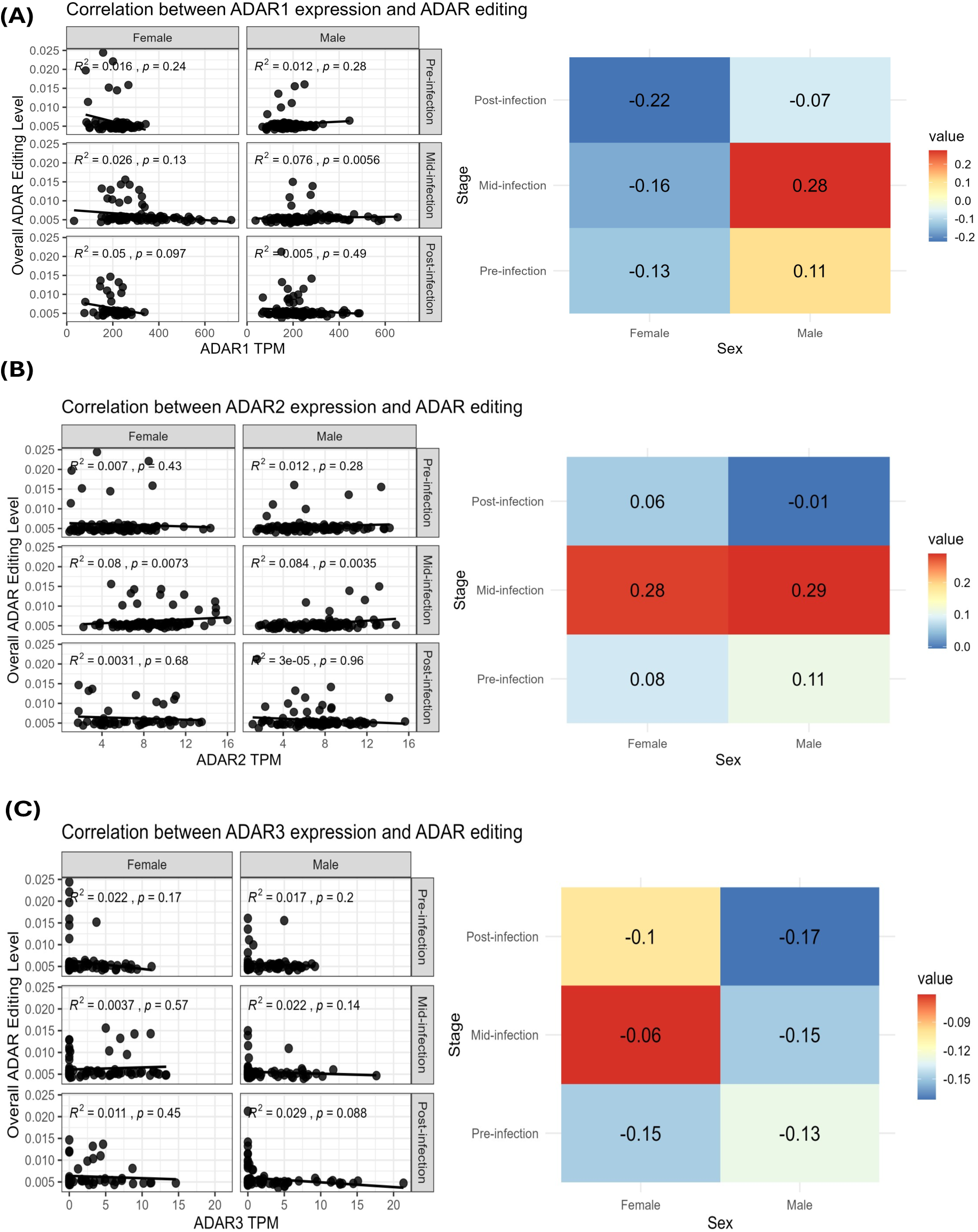
Correlations between the expression levels of all three ADARs (TPMs) and overall ADAR editing levels across the three stages of infection (pre-, mid-, and post-infection) in males and females. Each scatter plot shows the ADAR expression (on X axis) versus overall ADAR editing levels (on Y axis). Colored heatmap displays Spearman rank correlation coefficients (*ρ)* values and the color scale of the heatmap indicates the magnitude and direction of correlation. (A) Correlation between ADAR1 expression and overall ADAR editing shows that editing levels are significantly and positively correlated with ADAR1 expression only in males, mid-infection (*ρ =* 0.28, R^2^ = 0.076, p = 0.0056). (B) Displays significant correlation between ADAR2 expression and ADAR editing in both males (*ρ =* 0.29, R^2^ = 0.084, p = 0.0035) and females (*ρ =* 0.028, R^2^ = 0.08, p = 0.0073), mid-infection. While editing levels, pre- and post-stages of infection do not display significant correlations to ADAR2 expression (C) editing levels are not correlated with ADAR3 expression across any comparison group.

### ADAR editing within X-chromosome inactivation (XCI) genes

XCI is a process in females where one of the two X chromosomes is silenced to balance gene expression between the sexes. However, XCI is incomplete in humans, and approximately 23% of the X-chromosomal genes escape inactivation, which results in sex-specific gene expression patterns and phenotypic diversity. Many of innate immune related genes are encoded on the X-chromosome and their differential expression is associated with differential responses and manifestations during viral infections (Posynick & Brown, 2019; Tukiainen et al., 2017).

Therefore, using a curated list of genes that escape XCI from Tukiainen et al. study (Tukiainen et al., 2017), we investigated whether ADAR-mediated editing of genes that escape XCI display sex-specific differences across stage of SARS-CoV-2 infection as a potential post-transcriptional mechanism contributing to differential host responses between males and females. Weighted ADAR editing levels within XCI escape genes displayed significant variations between the sexes across different stages of infection. In pre-infection there were significant differences in weighted editing levels, with males exhibiting higher weighted editing compared to females, (Wilcoxon rank-sum test: p-value = 0.0059, Cliff’s δ = -0.237) (Supplementary Table 3 A, B) (Fig 4A).

**Figure 4:**
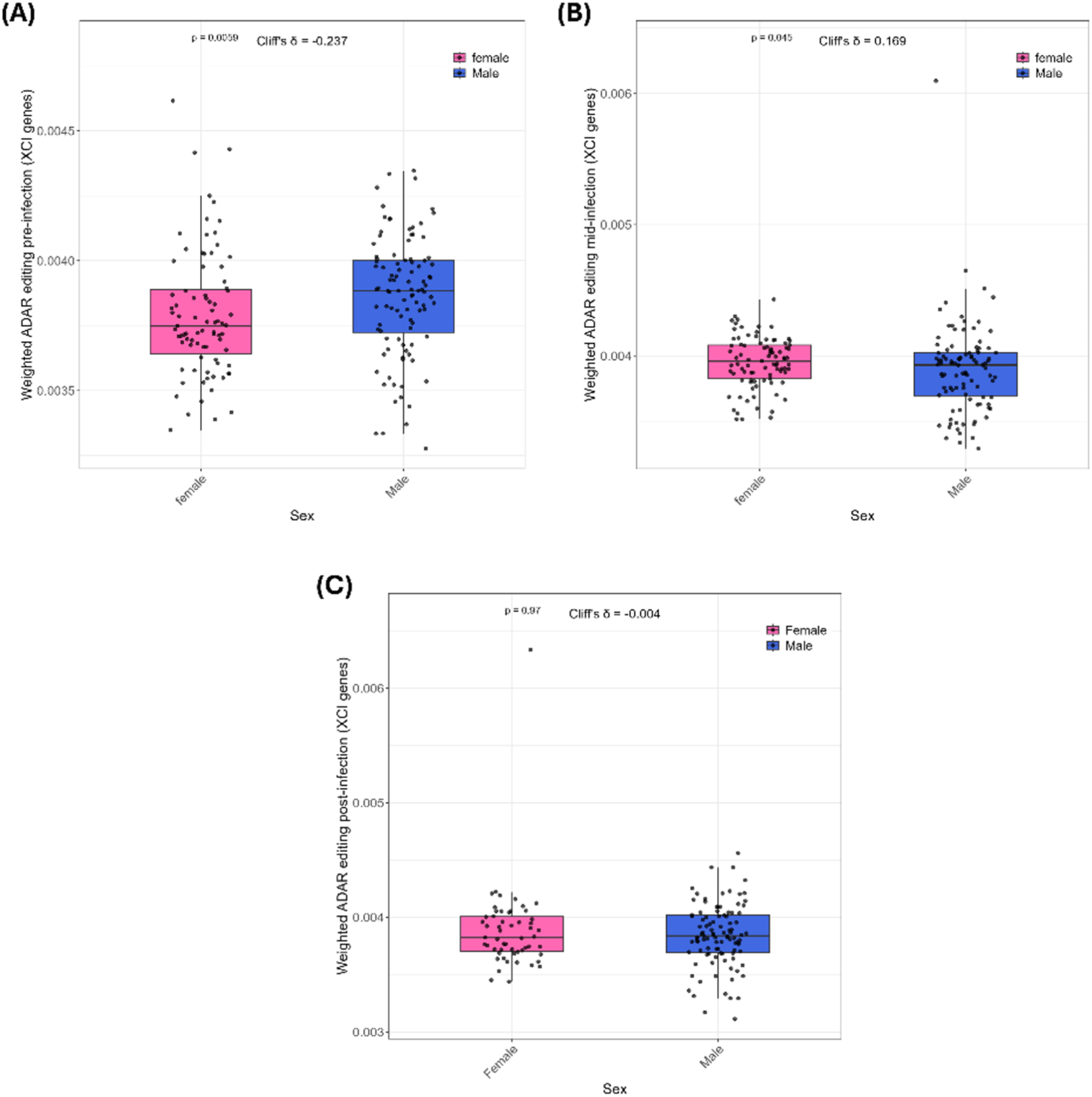
Sex-specific differences in weighted ADAR editing of XCI escape genes. (A) Pre-infection displayed statistically significant differences in ADAR editing levels between sexes (p-value= 0,0059), with males exhibiting slightly higher editing (Cliff’s δ = -0.237). (B) Mid infection editing shifted with females displaying significantly higher editing (p-value = 0.045), though effect size is low (Cliff’s δ = 0.169). (C) post infection analysis showed no significant differences (p-value = 0.97, Cliff’s δ = -0.004).

Interestingly, this difference reversed mid-infection, with females showing significantly higher editing levels than males (Wilcoxon rank-sum test: p-value = 0.045, Cliff’s δ = 0.169) (Supplementary Table 3 C, D) (Fig 4B). By post-infection there were no significant differences in weighted editing levels between the sexes (Wilcoxon rank-sum test: p-value = 0.0.97, Cliff’s δ = -0.004) (Supplementary Table 3 E, F) (Fig 4C). These results demonstrate that ADAR editing within XCI escape genes are dynamically modulated in a sex-dependent manner according to the stages of infection. While the observed effect size is small, it is important to note that ADAR-mediated RNA editing is a highly nuanced regulatory process and that even a small change can have profound biological impacts downstream (Watanabe et al., 2014).

## Discussion

Since the beginning of the COVID-19 pandemic sex has been identified as an important variable affecting COVID-19 outcomes (Takahashi et al., 2020). ADARs are RNA editing enzymes whose expression is altered during viral infections resulting in changes to host transcriptomic ADAR editing patterns (Piontkivska et al., 2021; Tariq & Piontkivska, 2024). These alterations to normal editing patterns could result in physiological changes during disease states (Slotkin & Nishikura, 2013). Differential expression of IFNs significantly contributes to differences in outcomes observed between males and females during SARS-CoV-2 infection (Takahashi et al., 2020). Considering that ADARs are downstream effector genes of the IFN pathway, here we explored sex-specific differences in ADAR levels and editing patterns between males and females, in pre-, mid-, and post-stages of SARS-CoV-2 infection. Our results demonstrate substantial differences in expression of all three ADAR genes, and global editing measured as differences in the total number of edits and the overall editing rates. We further reveal the dependence of global editing on expression of ADARs, and potential differences in activity of ADARs between the two sexes. Furthermore, we identified dynamic RNA editing within XCI escape genes as a previously underappreciated molecular mechanism that may contribute to sex-specific immune regulation and differential disease manifestation during COVID-19.

One of the strengths of our study is the availability of pre-infection samples from the same participants, allowing us to analyze baseline differences in expression and editing of ADARs between the sexes. In our samples, pre-infection ADAR1 levels were significantly higher, while ADAR2 levels were significantly lower in females. Noteworthy, this difference in expression of catalytically active ADARs was not reflected downstream in global editing levels. This apparent discrepancy could be due to compensatory shifts within the edited sites. ADAR1 and ADAR2 have overlapping as well as distinct targets and often compete for overlapping targets (Costa Cruz et al., 2020; Riedmann et al., 2008). The higher-expressed ADAR1 in our samples could thus outcompete lower-expressed ADAR2 at shared sites, resulting in higher and lower levels of ADAR1- and ADAR2-driven editing, respectively. Likewise, global editing measures may not show differences, because infection-driven gain in editing within some sites might cancel out the loss in others. If this holds true, at baseline the differences in editing might be qualitative (i.e., which specific site is edited) rather than quantitative. This hypothesis is supported by the presence of edited sites uniquely edited between males and females pre-infection, similar to the shift in edited sites observed in Reovirus infection (Tariq & Piontkivska, 2024). However, future studies using ADAR1 and ADAR2 specific knockdown approaches are needed to test whether (and how) inverse relationship between ADAR1 and ADAR2 expression results in redistribution of edited sites without a detectable net change in global editing.

Interestingly, in mid-infection, levels of both catalytically active ADAR1 and ADAR2 genes were elevated. Consistent with this, we observed higher overall ADAR editing levels in females at this stage. However, correlation analysis indicates that the relationship between ADAR expression and global editing differs by sex and stage of infection. For instance, during infection, ADAR2 levels were significantly and positively correlated with the overall ADAR editing only mid-infection. Similarly, ADAR1 correlated with global editing levels in males but not in females, despite higher ADAR1 expression and increased levels of editing within females mid-infection. These findings suggest that the contribution of ADAR levels to global editing varies according to sex-stage contexts, and that additional regulatory factors beyond the abundance of ADARs may influence global editing rates. Notably, expression of ADAR3, the catalytically inactive ADAR (Karlström et al., 2024), was also elevated in females during infection. ADAR3 expression has been reported to be largely restricted to the brain (Karlström et al., 2024), where it is thought to inhibit the activity of the other two ADARs. Although few studies have identified negligible ADAR3 expression in other organs such as liver and in gonadal tissues (Heraud-Farlow et al., 2017), its functional roles, if any, remain poorly understood. While ADAR3 expression was negligible in males mid-infection, its potential functional relevance of elevated expression in female warrants further attention.

We also observed sex-specific and stage-dependent differences in editing levels within genes that escape XCI. Notably, despite the absence of significant differences in global editing between the sexes at pre-infection stage, editing levels within XCI escape genes were significantly higher in males. This is intriguing, considering that higher expression of XCI escape genes (because they are expressed from both X chromosomes) would be expected to increase the number of potential ADAR-editable substrates and, consequently, influence their editing levels. Our finding suggests that RNA editing within XCI genes is not simply driven by the availability of substrates. Though the exact mechanism underlying this difference is unknown, a potential explanation is that ADAR-mediated editing could be a compensatory post-transcriptional regulatory mechanism to regulate expression of these genes in males. While in females, where there are two alleles being expressed for these genes, their expression may be regulated more at the transcriptional than post-transcriptional level. In contrast, the significant increase in editing within females during infection may be consistent with higher expression of ADARs at this stage. Given that many XCI escape genes play a role in immune regulation, increased editing within these genes may provide finer control of antiviral immune signaling in females. There were no sex differences post-infection within the periods of sample collection in the dataset, however, future studies with longer post-infection period would be required to identify if the levels go back to pre-infection states. These results imply difference in RNA editing within XCI escape genes as a potential additional layer of sex-specific immune regulation that could result in differential COVID-19 manifestations between sexes.

## Conclusion

Our study demonstrates dynamic and sex-specific differences in ADAR editing over the course of SARS-CoV-2 infection. By taking advantage of samples across pre-, mid-, and post-infection, we showed that differences in ADAR expression, the overall ADAR editing levels, and correlations between expression of ADARs and editing activity vary between males and females and that the editing is dynamically regulated over the course of infection. We also show that the sex differences are not merely because of change in ADAR expression levels, but could also be influenced by other mechanisms that regulate ADAR editing activity. Furthermore, in our analysis of RNA editing within genes that escape XCI, a gene set that has significant contribution to sex-specific outcomes to viral infections, we show that the effects of post-transcriptional ADAR editing, could be independent of gene expression. The dynamically changing editing landscape in these genes suggest that ADAR-mediated RNA editing might be an additional regulatory layer regulating immune related gene expression between males and females. Taken together, our results show ADAR-mediated RNA editing might contribute to differences in host responses and, to some extent, may explain differences in clinical COVID-19 manifestations between males and females.

## List of abbreviations

ACE2: angiotensin converting enzyme 2
ADAR: Adenosine Deaminase Acting on RNA
COVID-19: Coronavirus disease 2019
dsRNA: double stranded RNA
IFN: Interferon
ISG: Interferon stimulated gene
MERS: Middle East Respiratory Syndrome
SARS-CoV-2: Severe acute respiratory syndrome coronavirus 2
TMPRSS2: Transmembrane Serine Protease 2
XCI: X Chromosome Inactivation

## Institutional Review Board Statement

As a secondary analysis of publicly available, unrestricted access, data, this study was determined exempt human subjects research under the Category 4 of the Common Rule (45 CFR 46) by the Kent State University Institutional Review Board (protocol 2553).

## Informed Consent Statement

As a secondary analysis of publicly available, unrestricted access, data, this study is exempt from the consent requirements, under the Category 4 of the Common Rule (45 CFR 46).

## Data Availability Statement

The dataset used in this study is publicly available in the NCBI SRA/BioProject repository, as Bioproject PRJNA815324 and GEO accession GSE198449 (Sauerwald et al., 2022). Supplementary Materials and code are shared at https://github.com/RNAdetective/Sex-specific_differences_in_ADAR_editing_in_COVID19.

